# Strong preference for autaptic self-connectivity of neocortical PV interneurons entrains them to γ-oscillations

**DOI:** 10.1101/477554

**Authors:** Charlotte Deleuze, Gary S. Bhumbra, Antonio Pazienti, Caroline Mailhes, Andrea Aguirre, Marco Beato, Alberto Bacci

**Author notes:** Co-senior and co-corresponding authors.

## Abstract

Parvalbumin (PV) positive interneurons modulate cortical activity through highly specialized connectivity patterns onto excitatory pyramidal neurons (PNs) and other inhibitory cells. PV cells are auto-connected through powerful autapses, but the contribution of this form of fast disinhibition to cortical function is unknown. We found that autaptic transmission represents the most powerful input of PV cells in neocortical Layer V. Autaptic strength was greater than synaptic strength onto PNs as result of a larger quantal size, whereas autaptic and heterosynaptic PV-PV synapses differed in the number of release sites. Overall, single-axon autaptic transmission contributed to ~40% of the total perisomatic inhibition that PV interneurons received. The strength of autaptic transmission modulated the coupling of PV-cell firing with optogenetically-induced γ-oscillations preventing high frequency bursts of spikes. Autaptic self-inhibition represents an exceptionally large and fast disinhibitory mechanism to synchronize the output of PV cells during cognitive-relevant cortical network activity.

## Introduction

In the neocortex, cognitive-relevant processes depend on the activity of intricate networks formed by specific excitatory and inhibitory neuronal populations that are inter-connected according to a detailed blueprint (Allene et al., 2015;Harris and Shepherd, 2015;Kepecs and Fishell, 2014;Tremblay et al., 2016). In particular, fast synaptic inhibition governs both spontaneous and sensory-evoked cortical activity, and originates from a rich diversity of cell types with precisely distinct functions within cortical circuits (Isaacson and Scanziani, 2011;Tremblay et al., 2016). Perisomatic-targeting parvalbumin (PV)-expressing basket cells represent a major population of cortical GABAergic neurons. By providing fast inhibition onto PN cell bodies, PV cells exert a fine control of their output gain (Atallah et al., 2012;Tremblay et al., 2016) and spike timing, resulting in the generation and modulation of γ-rhythms, important for sensory perception and attention (Bartos et al., 2007;Buzsaki and Wang, 2012;Cardin et al., 2009;Sohal et al., 2009). Indeed, in awake animals, PV cells fire trains of spikes, which are strongly phase-locked to both spontaneous and visually evoked γ-rhythmic activity (Perrenoud et al., 2016).

In addition to targeting PNs, PV cells strongly inhibit one another, and GABAergic connections between PV cells contribute the greatest inhibition of this interneuron type (Avermann et al., 2012;Pfeffer et al., 2013). Moreover, PV cells are self-connected by autapses (synapses that a neuron makes with itself (Van der Loos and Glaser, 1972)). Self-inhibition was first described anatomically in adult neocortex of the cat (Tamas et al., 1997), and it was demonstrated to be functional in rodent (Bacci et al., 2003;Bacci and Huguenard, 2006;Connelly and Lees, 2010;Deleuze et al., 2014;Manseau et al., 2010) and human neocortex (Jiang et al., 2012;Jiang et al., 2013). In particular, fast autaptic neurotransmission plays a crucial role in setting millisecond-precise spike timing of PV cells during trains of action potentials (Bacci and Huguenard, 2006). Moreover, high-frequency firing of PV cells triggers massive asynchronous autaptic release of GABA, resulting in prolonged PV-cell self-inhibition that desynchronizes PV cell firing (Jiang et al., 2013;Manseau et al., 2010) (Jiang et al., 2012).

Connections between PV cells form a specific inhibitory network that is important for synchronizing a large population of neurons during γ-oscillations (Bartos et al., 2007;Buzsaki and Silva, 2012;Buzsaki and Wang, 2012). Despite the known role of PV cells as the clockwork of cortical networks, the underlying mechanism is still poorly understood. In addition, although functional autaptic transmission was demonstrated, the actual proportion of self-connections in relation to other synaptic projections from neocortical PV cells to other cells is unknown. Are autapses solely a connectivity curiosity, or do they represent an important source of inhibition of PV cells? Could fast self-inhibition contribute in keeping PV cell firing in sync with rhythmic network activity?

Here we measured the strength of autaptic self-inhibition compared to synaptic transmission from the same PV cell onto their two principal synaptic targets: PNs and other PV cells. Remarkably, autaptic responses were invariably much larger than unitary synaptic transmission onto PNs, and, on average, onto other PV cells. Quantal parameters underlying the autaptic vs. synaptic strength were different depending on whether the postsynaptic neuron was a PN or another PV cell. We found that PV cells with strong autaptic inhibition provided little input to other PV cells, while PV cells with smaller autapses provided larger heterosynaptic inhibition to neighboring PV cells. Remarkably, self-connections accounted for up to ~40% of the entire inhibitory strength onto single PV cells. Finally, we found that autaptic transmission tuned the strong coupling of PV-cell spikes with γ-oscillations, by modulating spike after-hyperpolarization (AHP) and thus inter-spike intervals. Therefore, autaptic self-innervation accounts for a large fraction of synaptic inhibition PV cells receive, and is responsible for locking their spiking activity to cognitive-relevant network oscillations.

## Results

### Layer V PV cells connect more powerfully with themselves via autaptic contacts than with other synaptic partners

In order to compare autaptic inhibition of PVs cells with the synaptic inhibition from the same PV cells through GABAergic connections onto PNs and other PV cells, we performed simultaneous paired recordings between these two cell types in neocortical Layer V of the mouse somatosensory (barrel) cortex of acute brain slices. We used a transgenic PV-cre∷tdTomato mouse strain to identify PV cells unambiguously (see Methods). Briefly, tdTomato-positive neurons exhibit a clear multipolar, aspiny morphology and stereotypical fast-spiking behavior. Non-fluorescent PNs had typical large cell bodies and an apical dendrite directed towards the pia (see Methods). We isolated GABAergic events pharmacologically, and used a high-Cl intracellular solution (see Methods) for voltage-clamp recordings of GABAergic synaptic currents that were inward at a holding potential of −80 mV. We elicited action currents in PV cells in voltage-clamp by delivering brief (0.2 - 0.6 ms) depolarizing steps from −80 mV to membrane potential between −20 mV to 0 mV in order to minimize passive electrical artefacts induced by the stimulus. Self-connected PV cells exhibited large GABAergic inward responses following action currents. As previously demonstrated (Bacci et al., 2003), this response results from unitary autaptic transmission, since it exhibits fixed latencies, peak amplitude functions, and were abolished by the GABA_A_R antagonist gabazine (10 µM, Fig. 1A). We found GABAergic autaptic inhibitory postsynaptic currents (autIPSCs) in 74% of recorded PV neurons (n = 164). The same action currents in PV cells elicited unitary inhibitory postsynaptic currents (synIPSCs) onto a fraction of PNs (Fig. 1A,B) or PV cells (Fig. 1C,D). The yield of finding connected PV-PN pairs was of 61% (36 out of 59 pairs), of which 75% exhibited also autaptic responses (27 out of 36 pairs). We found that PV-PN responses were invariably much smaller than their autaptic counterparts, either when they were analyzed independently (Table 1; p = 5.6E-7, n = 84 and 22 for autaptic and synaptic transmission, respectively), or when paired dual connections were analyzed separately (Table 2; p = 5.3E-4, n = 16; Fig. 1B). Among pairs between PV cells, the proportion of connected synaptic PV-PV pairs was 61% (59 out of 96 pairs), of which 76% exhibited also autaptic responses (45 out of 59 pairs). Also in this case, PV-cell autaptic strength was larger than PV-PV synaptic transmission (Table 1; p = 5.7E-5, n = 84 and 49 for autaptic and synaptic transmission, respectively), both when autIPSCs and synIPSCs were analyzed independently, and when paired dual connections were analyzed separately (Table 2; p = 0.0215, n = 38; Fig. 1D).

**Table 1:**
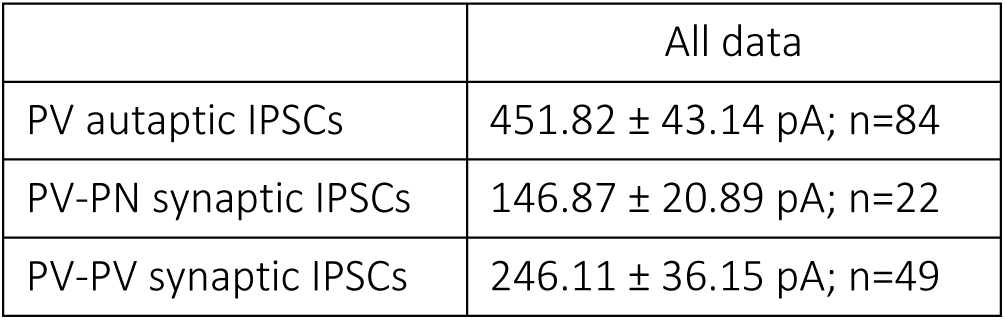
Mean current in all PV-PN and PV-PV pairs

|  | All data |
| --- | --- |
| PV autaptic IPSCs | 451.82 ± 43.14 pA; n=84 |
| PV-PN synaptic IPSCs | 146.87 ± 20.89 pA; n=22 |
| PV-PV synaptic IPSCs | 246.11 ± 36.15 pA; n=49 |

**Table 2:**
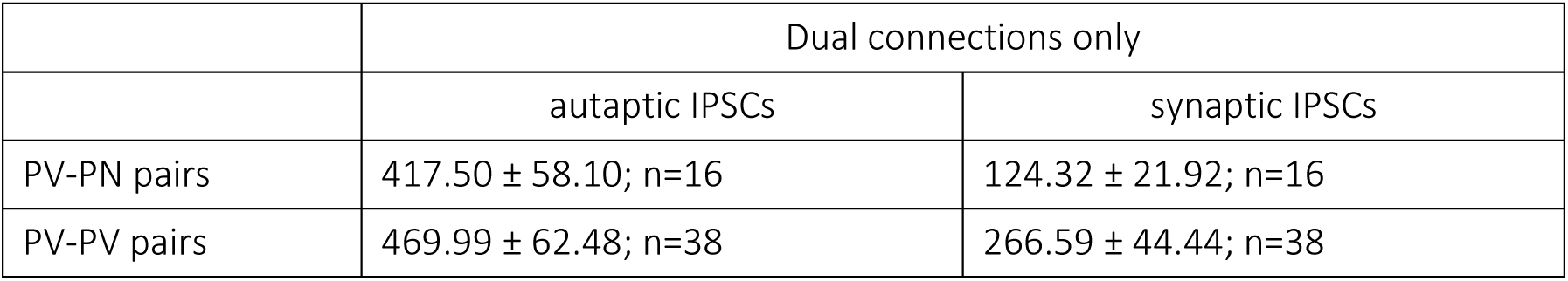
Mean current in PV-PN and PV-PV pairs with both autaptic and synaptic connections

**Figure 1:**
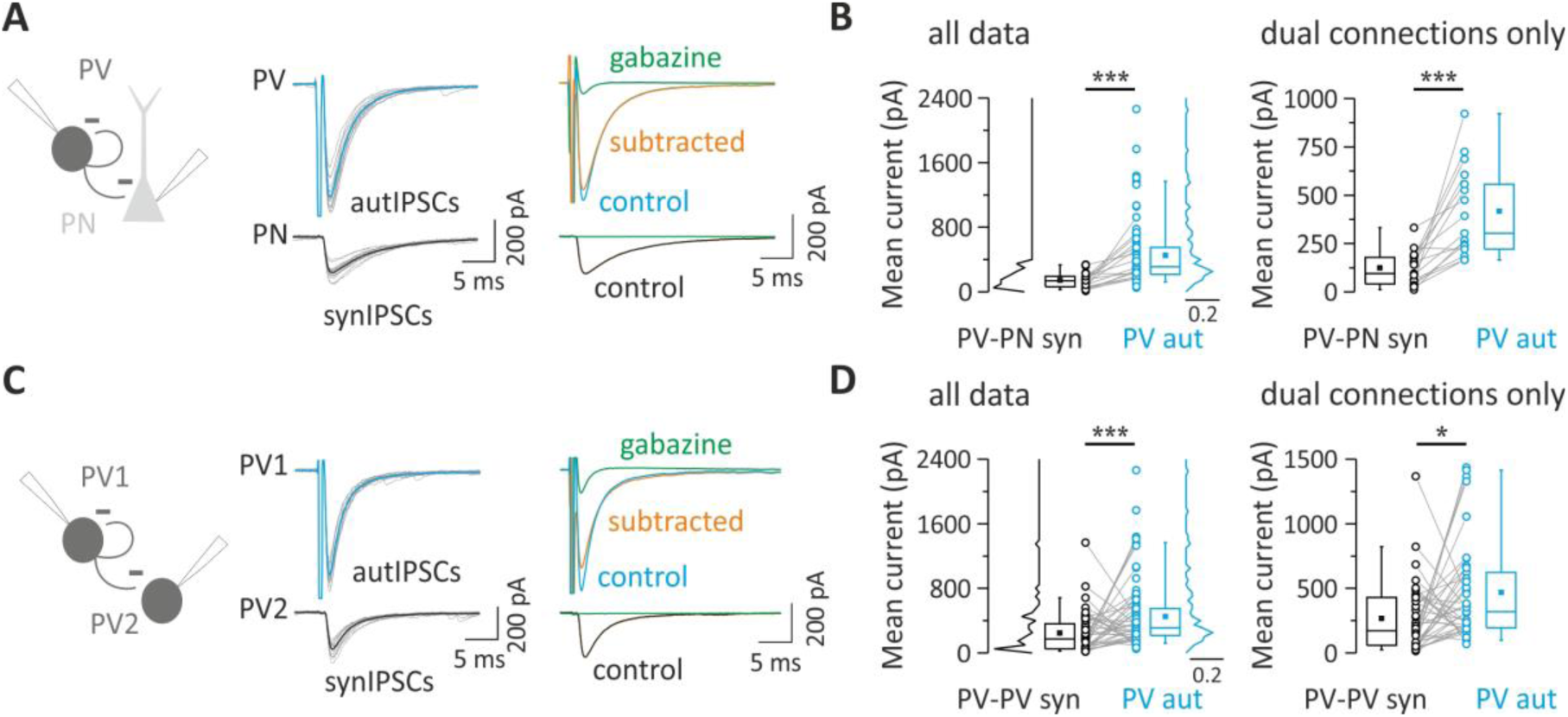
Layer V PV cells connect more powerfully with themselves via autaptic contacts than with other synaptic partners. ***A***, Unitary autaptic and synaptic inhibitory currents (autIPSCs and synIPSCs) evoked simultaneously in a PV cell and a PN respectively, in response to PV cell stimulation. Individual responses (15 grey traces) were averaged (thick trace, blue for autIPSC and black for synIPSC). In the presence of the GABA_A_R antagonist, gabazine (10 µM), the two responses were blocked but note the residual current in the PV cell reflecting the distortion due to the voltage step eliciting the action potential current (clipped). In order to cancel this stimulus waveform, current traces in gabazine were subtracted from control responses (orange: subtracted trace, average of 10 trials). ***B***, Population data obtained from PV-PN pairs with either single (autaptic or synaptic) or paired dual (autaptic and synaptic) connections (all data, left panel). Right panel illustrates exclusively pairs with both synaptic and autaptic connections from the same presynaptic PV cell (dual connections only). Note that the mean autaptic current from PV cell is systematically and significantly larger than the synaptic one (*** p<0.001). ***C***, Representative traces of autIPSCs and synIPSCs as in A, but recorded in a PV-PV pair. ***D***, Population data obtained from PV-PV pairs with summary plots as described in B. Note that on average autaptic currents are larger than synaptic currents ((*** p<0.001, * p<0.05).

These results indicate that autaptic self-inhibition of PV cells is more powerful than synaptic transmission from the same PV cells onto their principal post-synaptic targets in Layer V: PNs and other PV cells.

### Quantal parameters accounting for larger unitary autaptic than synaptic connections between PV cells and PNs

Synaptic efficacy results from the combination of pre- and postsynaptic factors, namely the number of presynaptic release sites (n), the probability of neurotransmitter release (P_r_), and the postsynaptic response to a single released synaptic vesicle (or quantum), known as quantal size (q). This can be summarized by expressing the average unitary synaptic current <I_syn_> as a product of the quantal parameters:

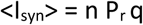

Autaptic self-inhibition onto PV cells is powerful and, on average, stronger than synaptic transmission from the same cells to other elements of cortical microcircuit (Fig. 1). We therefore set out to determine the quantal parameter(s), responsible for stronger autaptic neurotransmission using Bayesian quantal analysis (BQA) (Bhumbra and Beato, 2013). We recorded from pairs of PV cells and PNs, exhibiting GABAergic autaptic and synaptic responses, at two extracellular Ca^2+^ concentrations (2.0 and 1.5 mM), resulting in different release probabilities. At the end of each recording, we applied the GABA_A_R antagonist gabazine to enable subtraction of the stimulus waveform and action current to isolate autaptic responses for each of the Ca^2+^ concentrations (Fig 2A). Also in this set of data, autIPSCs recorded with high ([Ca^2+^] (2 mM) were invariably larger than synIPSCS (mean current = 394.15 ± 58.54 vs. 151.64 ± 26.24 pA; autaptic vs. synaptic connections; p = 0.002, n = 11; Fig. 2B,C). We then applied BQA at unitary autaptic and synaptic responses recorded at low ([Ca^2+^] = 1.5 mM) and high ([Ca^2+^] = 2 mM) release probabilities, and obtained median-based estimates for the quantal parameters from the marginal posterior distributions for the quantal size q and maximal response r (where r = nq), and hence number of release sites n (Fig. 2 D, E) (Bhumbra and Beato, 2013).

**Figure 2:**
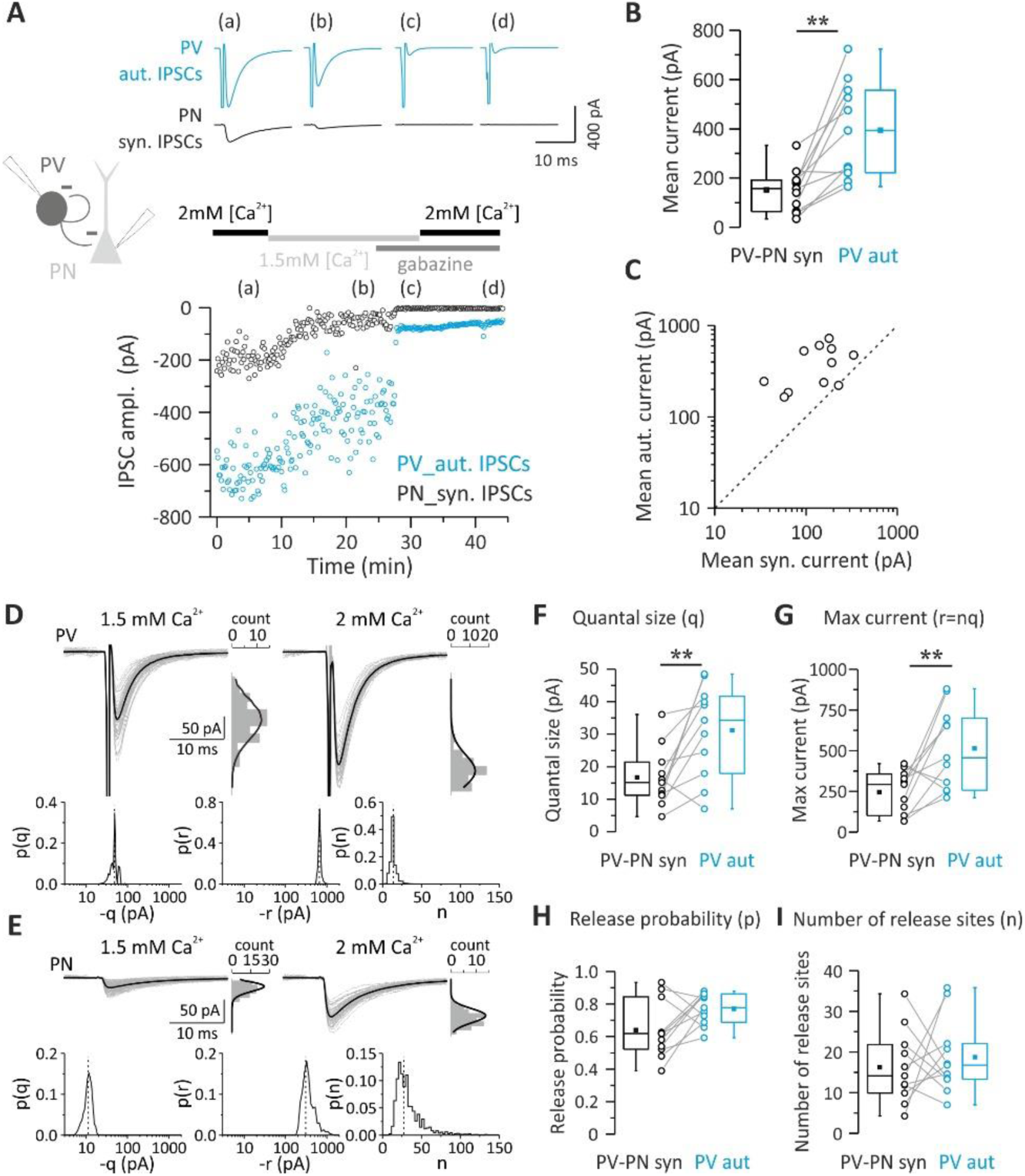
Quantal parameters accounting for larger unitary autaptic than synaptic connections between PV cells and PNs. ***A***, Top: Representative autaptic and synaptic traces in response to PV-cell stimulation recorded from a PV-PN pair. IPSCs were elicited every 10 sec and each trace is the average of 10 sweeps. Bottom: time course of autIPSC (blue symbols) and synIPSC (black symbols) amplitudes recorded simultaneously from the same PV cell and PN, respectively, at two extracellular Ca^2+^ concentrations (2.0 and 1.5 mM) and in the presence of 10 µM gabazine (at both [Ca^2+^]) to subtract the stimulus waveform. ***B-C***, Summary (B) and correlation plots (C) of synIPSCs and autIPSC amplitude obtained from all PV-PN pairs with dual connections used for BQA and measured at 2 mM [Ca^2+^]. Note that in all pairs (open circles), autaptic currents in PV cells are consistently bigger than their synaptic correlates in PN as all points fall above the unity line (dashed line). ***D-I***, Quantal analysis of PV and PN responses to PV-cell stimulation recorded from the PV-PN pair shown in A. Example responses of the PV cell (***D***) and PN (***E***) are shown alongside their respective amplitude distributions observed in the presence of of 1.5 mM (left, low release probability) and 2 mM (right, high release probability) extracellular Ca^2+^. The results of BQA are represented as probability distributions for the quantal size (***F***; q), maximal response (***G***, n), release probability (***H***, p) and number of release sites (***I***; r). The dashed line is the median. Note the larger quantal size and maximum current of autaptic responses. (n=11; ** p<0.01).

We found that in PV-PN pairs with both autaptic and synaptic connections, autaptic responses had a significant larger quantal size (q) than unitary synaptic connections onto PNs (Table 3; p = 0.00976, n = 11; Fig. 2F). Accordingly, the maximal response r (nq) was also larger in autaptic vs. synaptic responses onto PNs (Table 3; p = 0.0098, n = 11; Fig. 2G). Conversely, no differences in release probability (Pr) and number of release sites (n) were shown by comparison of autaptic transmission onto PV cells and synaptic inhibition from the same neurons onto PNs (Table 3; p>0.05 n = 11; Fig. 2H-I).

**Table 3:** Bayesian quantal analysis in PV-PN pairs with both autaptic and synaptic connections

|  | autaptic IPSCs (n=11) | synaptic IPSCs (n=11) |
| --- | --- | --- |
| Quantal size (q) | 31.17 ± 4.28 pA | 16.68 ± 2.71 pA |
| Number of release site (n) | 18.70 ± 2.76 | 16.26 ± 2.70 |
| Probability of release (p) | 0.77 ± 0.03 | 0.64 ± 0.05 |
| Max current (r) | 513.65 ± 74.82 pA | 245.33 ± 39.88 pA |

These results indicate a larger quantal size at autaptic sites, as compared to synapses that the same PV cells formed with PNs. This is consistent with PV-PN synaptic transmission being invariably smaller than autaptic transmission. Therefore stronger autaptic efficacy is likely due to cell type-specific postsynaptic mechanisms.

### The strength of autaptic and synaptic transmission onto PV cells depends on different number of release sites

Pairs of PV cells were analyzed to determine whether differences in unitary autaptic vs. synaptic transmission onto other PV cells could be accounted for by any of the quantal parameters. We noticed that the strength of self-vs. heterosynaptic inhibition defined two connectivity patterns of PV cells: those with stronger autaptic than synaptic PV-PV connections, and those, which showed an opposite trend (referred to as ‘introverted’ and ‘extroverted’ PV cells, respectively; Fig. 3A,B). In our hands, ‘introverted’ PV cells (in which autIPSCs > synIPSCs) corresponded to 63.1% of the total dual connected sample (n = 24 out of 38). Of those 38 PV cells, stable experiments suitable for BQA analysis were obtained in 15 pairs, 9 of which were ‘introverted’ and the remaining 6 ‘extroverted’ PV cells (corresponding to 60 nd 40%, respectively; Fig. 3C,D). Importantly, the size of autaptic and unitary synaptic response were inversely correlated (R = −0.5643, p=0.031; Fig. 3D), suggesting the existence of two connectivity patterns between PV cells that could be distinguished by their self-inhibition strength. We found that in both ‘introverted’ and ‘extroverted’ PV cells, autaptic and synaptic quantal size (q) was similar for both ‘introverted’ and ‘extroverted’ PV cells (n = 9 and 6, respectively; Fig. 3E). In addition, release probability (Pr) was similar for self- and PV-PV synaptic inhibitory contacts; Fig 3F). However, we found that a differential number of release sites (n) determined the strength of autaptic and synaptic connections onto PV cells. Indeed, in 8 out of 9 ‘introverted’ PV-cell pairs, the number of release sites (n) was larger in autaptic than synaptic connections (Table 4; Fig. 3G, red symbols). Accordingly, the opposite was true for ‘extroverted’ PV-cell pairs, in which in 5 out of 6 cases, the number of autaptic release sites was smaller than PV-PV synaptic connections (Table 4; Fig. 3G, grey symbols). Therefore, the maximal autaptic response r (or nq) was larger or smaller in ‘introverted’ and ‘extroverted’ PV cell pairs, respectively (Table 4; Fig. 3H).

**Table 4:**
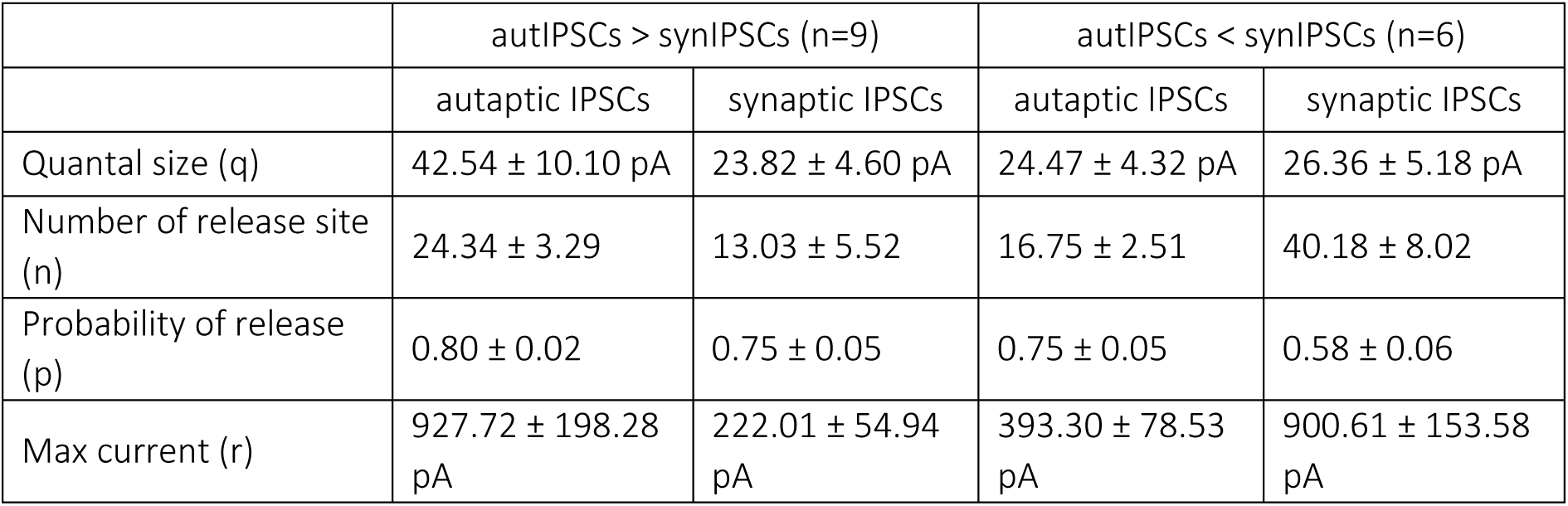
Bayesian quantal analysis in PV-PV pairs with both autaptic and synaptic connections

|  | autIPSCs > synIPSCs (n=9) |  | autIPSCs < synIPSCs (n=6) |  |
| --- | --- | --- | --- | --- |
|  | autaptic IPSCs | synaptic IPSCs | autaptic IPSCs | synaptic IPSCs |
| Quantal size (q) | 42.54 ± 10.10 pA | 23.82 ± 4.60 pA | 24.47 ± 4.32 pA | 26.36 ± 5.18 pA |
| Number of release site (n) | 24.34 ± 3.29 | 13.03 ± 5.52 | 16.75 ± 2.51 | 40.18 ± 8.02 |
| Probability of release (p) | 0.80 ± 0.02 | 0.75 ± 0.05 | 0.75 ± 0.05 | 0.58 ± 0.06 |
| Max current (r) | 927.72 ± 198.28 pA | 222.01 ± 54.94 pA | 393.30 ± 78.53 pA | 900.61 ± 153.58 pA |

**Figure 3:**
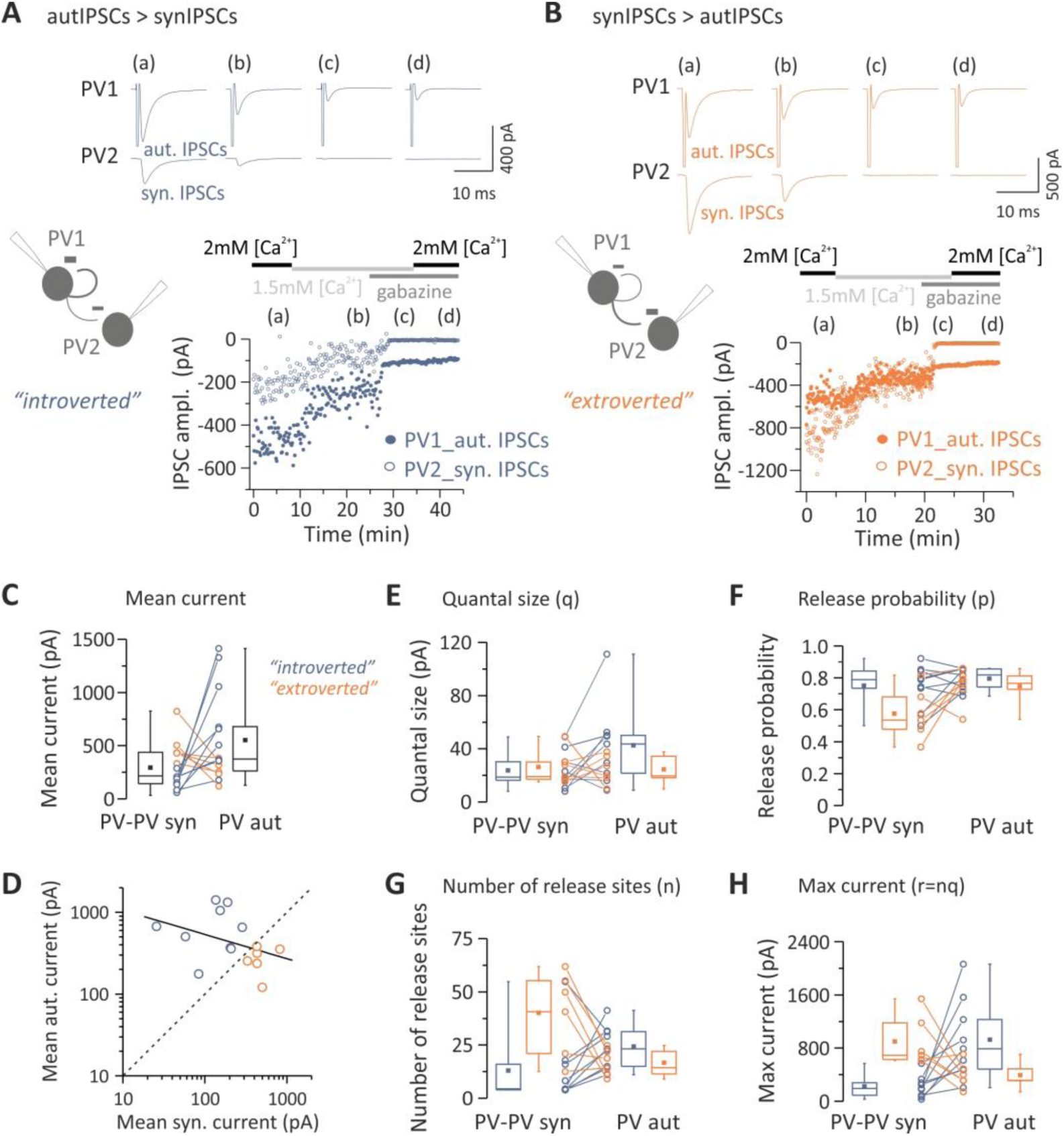
‘Extroverted’ and ‘introverted’ PV cells rely on a different number of release sites. ***A****-****B***, Top: Representative responses from pairs of PV cells (PV1 and PV2) at two extracellular Ca^2+^ concentrations (2.0 and 1.5 mM) and in the presence of gabazine. Shown are cases of ‘introverted’ (autIPSCs>synIPSCs, A) and ‘extroverted’ (autIPSCs<synIPSCs, B) presynaptic PV cells. Each trace is the average of 10 sweeps. Bottom: Time course of autaptic and synaptic IPSC amplitudes. Responses were elicited every 10 (***A***) or 5 sec (***B***). ***C***, Summary plots of synIPSCs and autIPSC amplitude from all pairs used for BQA (black box charts), measured at 2 mM [Ca^2+^], with color-coded ‘introverted’ and ‘extroverted’ PV cells (blue and orange, respectively). ***D*,** autIPSCs plotted against synIPSCs. Data were fitted with a linear regression (black line) showing a good degree of correlation between paired autaptic and synaptic responses from a single presynaptic PV cell‥ Note also that individual pairs (open circles) are distributed on both sides of the relationship for equal autaptic and synaptic IPSCs (dashed line), thus indicating a split of PV cells into two types of connection pattern, introverted and extroverted (blue and orange, respectively). ***E-H***, Quantal analysis of PV responses. ‘Introverted’ and ‘extroverted’ cases are color-coded as in A. Results of BQA are represented as probability distributions for the quantal size (***E***; q), release probability (***F***, p), number of release sites (***G***, n) and maximal response (***H***; r). Note that the large size of an autaptic/synaptic response relies on a large number of release sites compared to their synaptic/autaptic correlate in all but one introverted and extroverted PV cell, respectively.

These results indicate that the strength of autaptic transmission in PV cells, as compared to heterosynaptic PV-PV connections, is determined by the number of release sites and thus, can be accounted for by structural differences, in contrast with PV-PN connections, where the greater strength of autaptic vs. heterosynaptic currents is mainly determined by differences in the quantal size.

### Autaptic neurotransmission accounts for a large fraction of the total inhibition onto single PV cells

The prevalence of self-vs. synaptic GABAergic transmission originating from PV cells prompted the question of whether autaptic transmission provides a large proportion of the total synaptic inhibition that these cells receive. To measure the autaptic fraction contributing to the overall perisomatic inhibition received by single PV cells, we progressively blocked autaptic neurotransmission while evoking GABA release from virtually all terminals impinging the cell bodies of recorded PV cells. Autaptic transmission was blocked by intracellular perfusion of the fast Ca^2+^ chelator BAPTA (20 mM membrane impermeable free acid, in the presence of 2 mM Ca^2+^). Diffusion of BAPTA to autaptic contacts progressively reduced the size of autIPSCs until a complete block of autaptic neurotransmission was typically achieved within 20 minutes following whole-cell break in (Fig. 4A). In order to rule out that this time-dependent reduction of autaptic responses was due to non-specific rundown, we performed some control experiments, in which autIPSCs were recorded with an intracellular solution containing low concentration (1 mM) of the slow Ca^2+^ chelator EGTA, mimicking endogenous Ca^2+^ buffering by parvalbumin (Collin et al., 2005). In the presence of intracellular EGTA, autIPSCs were stable for long periods (up to 1 hour, Fig. 4B) (Bacci et al., 2003;Manseau et al., 2010).

**Figure 4:**
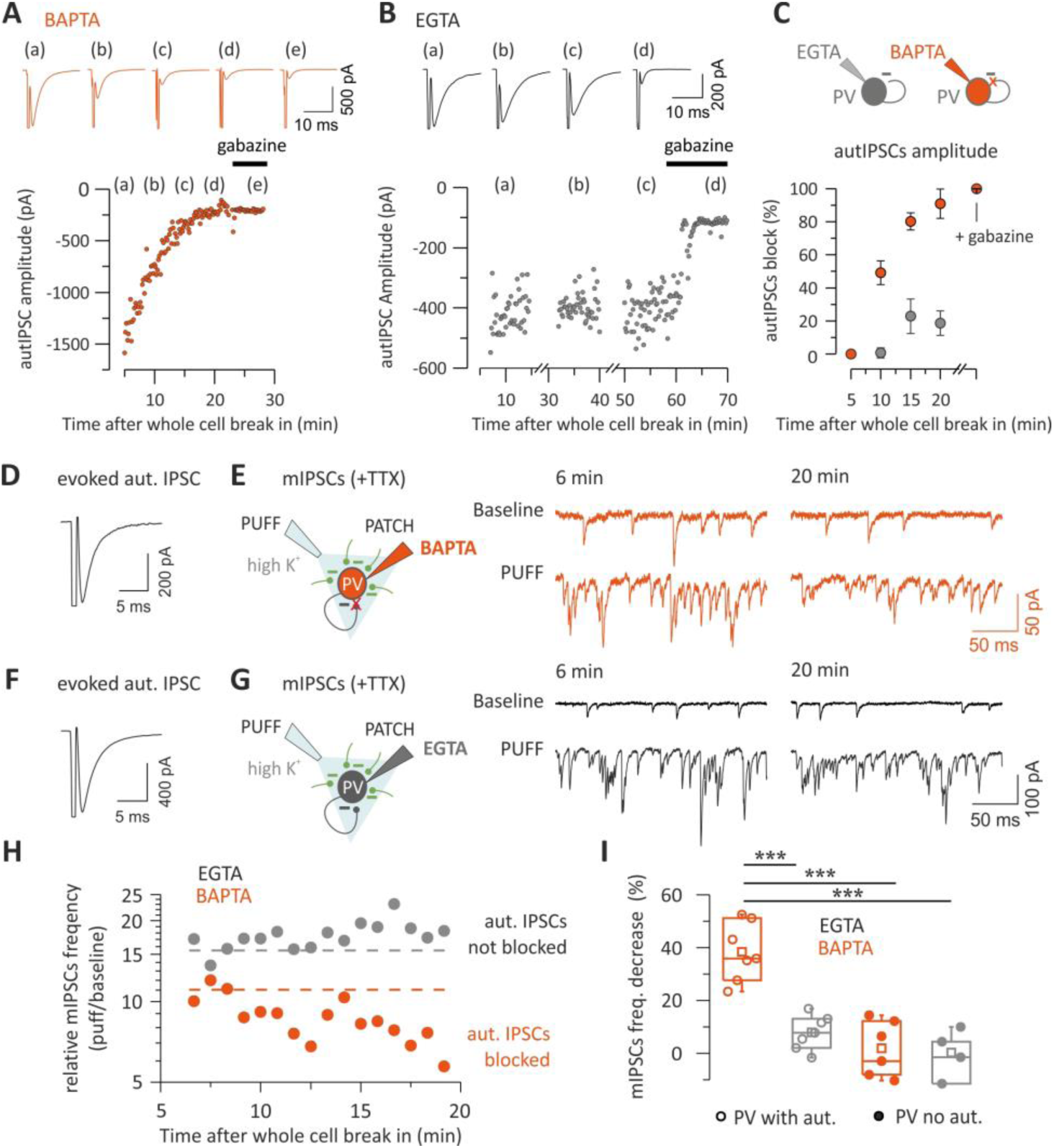
Autaptic neurotransmission accounts for a large fraction of the total inhibition onto single PV cells. ***A-C***, Representative voltage-clamp traces (top) and time course (bottom) of autaptic responses recorded from two PV cells in the presence of 20 mM BAPTA (A, orange) or 1 mM EGTA (B, black) in the recording whole-cell pipette. IPSCs were elicited every 10 sec and their amplitudes illustrated on the time course plot (bottom). Note the decline of autIPSC amplitudes during BAPTA perfusion up to a complete block (***A***), as compared to the absence of rundown during long (1 hr) EGTA perfusion (***B***). Gabazine completely blocked autIPSCs in both cases. ***C***, Summary of autaptic IPSC block by intracellular BAPTA in PV cells. IPSCs amplitudes were normalized to the average value obtained 5 min after establishment of whole-cell configuration. Autaptic currents are blocked by BAPTA perfusion within 20 min (n=5) but not in EGTA (n=5). ***D***, Average trace of autIPSC recorded in a PV cell immediately after whole-cell establishment, in the presence of BAPTA in the recording pipette. ***E,*** Schematic of the experiment (left) and representative voltage-clamp traces of mIPSCs before and after puffing high K^+^ ACSF (PUFF) at the beginning (6 min) and after intracellular blockade of autaptic release (20 min). Note the decrease of mIPSC frequency in high extracellular K^+^ after 20 minutes of intracellular perfusion of BAPTA. ***F-G***, Same as in E-F, but in another PV cell recorded with a control EGTA (1 mM) intracellular solution. Note that the increase of mIPSCs induced by high-K^+^ is constant over the same period. ***H***, mIPSCs frequency changes induced by high-K^+^ puff in the same PV cells as in ***E*** and ***G***, as estimated by the ratio before and after application of the high-K^+^ solution for each puff. The dashed line indicates baseline frequency (average of the 1^st^ three points). Note that in BAPTA, the frequency declines progressively following the same time course than the autaptic transmission block whereas it is stable in EGTA. ***I,*** The percentage of decrease in mIPSCs frequency calculated between the same time points as in E and G *i.e* before and after potential autaptic transmission block, is shown on the summary plot. In PV cells with evoked autaptic IPSC, the frequency strongly decreases in presence of intracellular BAPTA (n=7) but not EGTA (n=7) whereas in PV cells with no autaptic transmission, the frequency did not change in both conditions (BAPTA, n=6; EGTA, n=4) (*** p<0.001).

On average, after 20 min of intracellular BAPTA diffusion, autIPSC amplitude was 9.1 ± 8.8 % of control (n=5), whereas in the same timeframe, intracellular EGTA diffusion did not affect autaptic transmission (81.3 ± 7.4 % of control; n=5). To measure the relative fraction of autaptic inhibition onto PV cells, we first tested whether or not the recorded PV cell exhibited an autaptic response (Fig. 4D); we subsequently applied the Na^+^-channel blocker tetrodotoxin (TTX, 1 µM) and measured a baseline period of miniature mIPSCs. Using a local micropipette, we then puffed ACSF with a high concentration of KCl (20 mM) to depolarize all synaptic terminals impinging upon the recorded PV cell, thus forcing global Ca^2+^-dependent release of GABA without inducing unwanted network effects (Fig. 4E). We repeated the high K^+^ puffs once per minute, for at least 20 min, and we measured the relative increase of mIPSC frequency (puff/baseline) as an estimate of global perisomatic inhibition onto the recorded cell (Fig. 4 D-G).

In the presence of 20 mM intracellular BAPTA, the high-K^+^-dependent increase in mIPSC frequency declined steadily within 20 minutes after whole-cell break in (Fig. 4H), consistent with a complete autaptic blockade (Fig. 4B). In contrast, in control experiments in which EGTA was internally diffused, the increase of mIPSC frequency was stable over the same period of time (Fig. 4H). On average, mIPSC frequency blockade was 38.4 ± 4.2 % and 7.9 ± 2.4 % in the presence of BAPTA and EGTA, respectively (BAPTA n = 7; EGTA n=7; p < 3.9E-5; one way ANOVA, followed by Tukey’s comparison; Fig. 4I). Importantly, in those PV cells lacking autaptic responses, the high K^+^-dependent increase of mIPSC frequency was stable over time, regardless of whether intracellular BAPTA or EGTA was present, ruling out non-specific effects of BAPTA on mIPSCs. (1.9 ± 4.3 % and 0.3 ± 4.6 %; n = 6 and 4, BAPTA and EGTA, respectively; p = 0.99, one-way ANOVA; Fig. 4I).

These results indicate that, overall, unitary autaptic self-inhibition contributes to a large fraction (~40%) of the overall inhibition that PV cells receive.

### γ-Oscillations induced in Layers II/III are efficiently propagated to Layer V PV cells

PV cells play a key role in driving network oscillations in the β-γ-frequency range (20-100 Hz) (Bartos et al., 2007;Buzsaki and Wang, 2012;Cardin et al., 2009;Isaacson and Scanziani, 2011;Sohal et al., 2009), believed to underlie several cognitive functions, such as attention and sensory representation (Buzsaki and Silva, 2012;Isaacson and Scanziani, 2011). Importantly, PV-PV synaptic and electrical coupling is important for synchronizing these interneurons during γ-oscillations (Bartos et al., 2007;Buzsaki and Wang, 2012;Mann and Paulsen, 2007), and previous evidence indicated that autaptic self-inhibition of PV cells is instrumental for their spike precision in the γ-frequency range (Bacci and Huguenard, 2006). We therefore tested whether autapses, that constitute the major GABAergic output of these interneurons, could modulate PV-cell spike output induced by γ-like activity.

We expressed the light-sensitive opsin channelrhodopsin2 (ChR2) in a fraction of Layer II/III PNs via *in utero* electroporation of mouse embryos (Fig. 5 A,B; see Methods). ChR2-negative PNs in the same Layer II/III area were recorded where the opsin was expressed. In agreement with previous reports (Adesnik and Scanziani, 2010;Hakim et al., 2018;Shao et al., 2013), illumination of cortical slices with a ramp of blue light induced strong rhythmic activity of both IPSCs and EPSCs at ~30 Hz (Fig. 5C,D). Layer 2/3 PNs project monosynaptically to Layer V neurons (Adesnik and Scanziani, 2010). We therefore simultaneously recorded Layer V PV cells and ChR2-negative PNs in Layer II/III (Fig. 5E) to measure rhythmic IPSCs in Layer II/III PNs and voltage fluctuations of Layer V PV cells (see Methods). We found that light-evoked γ-activity in Layer II/III was reliably transmitted to Layer V PV cells, as shown by subthreshold PSPs, which oscillated at the same frequency of IPSCs recorded in Layer II/III (Fig. 5F,G). When the membrane potential was slightly depolarized, light activation of a fraction of Layer II/III PNs triggered several action potentials in Layer V PV cells (Fig. 5F), strongly resembling PV-cell firing activity recorded *in vivo (*Perrenoud et al., 2016).

**Figure 5:**
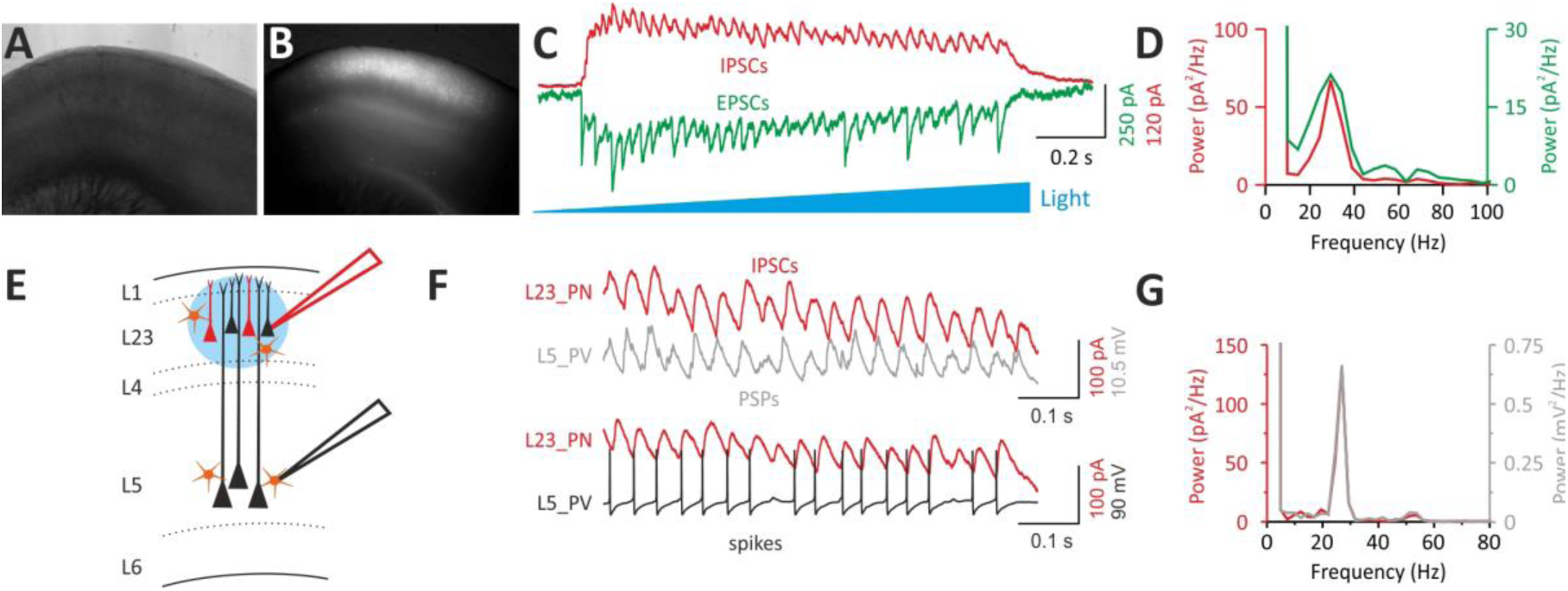
Optogenetically induced γ-oscillations in Layer II/III efficiently propagate to Layer V PV interneurons. ***A-B,*** Bright-field (A) and fluorescent (B) photomicrographs of an acute cortical slice of a mouse that was electroporated *in utero* at E15.5 with two plasmids expressing ChR2 and the red-fluorescent protein mRFP, respectively. Note the wide expression of mRFP in Layer II/III in the barrel field (B). ***C***, Representative voltage-clamp traces of IPSCs (red) and EPSCs (green) recorded from a ChR2-negative PN of layer 2/3 in response to a ramp of blue light delivered with an LED coupled to the epifluorescence path of the microscope. IPSCs and EPSCs were isolated by holding the recorded neuron at the reversal potential of glutamate- and GABA-mediated responses, respectively. ***D,*** Power spectra of the IPSCs (red) and EPSCs (green) of the cell of *C*. Note the sharp peak at ~30 Hz, in the γ-frequency range. ***E,*** Scheme of the experimental configuration: a dual patch-clamp recording is established. A ChR2-negative PN in layer 2/3 is recorded in voltage-clamp and a PV cell in Layer V is simultaneously recorded in current-clamp: a ramp of blue light is then delivered on Layer II/III PN cell bodies. ***F,*** A ramp of blue light induces rhythmic IPSCs in the Layer II/III PNs and subthreshold PSPs in the PV cell recorded at its resting potential in Layer V. When the PV cell was slightly depolarized, optogenetic activation of ChR2-positive Layer II/III PNs induced sustained firing of the PV cell in Layer V. ***G***, The power spectra of IPSCs (red) and PSPs (gray) of the Layer II/III PN and Layer V PV cell shown in *F* coincide, indicating a good transmission of Layer II/III γ-activity across the two cortical layers.

These results indicate that optogenetically induced γ-oscillations in Layer II/III are faithfully propagated to Layer V PV cells, thus allowing studying the role of autaptic self-innervation of these cells during cortical network activity.

### Autaptic neurotransmission is instrumental for locking PV-cell firing to γ-oscillations

We tested if the strong inhibitory autaptic conductance occurring after each spike in PV cells is important for synchronizing these interneurons during γ-oscillations. Autaptic responses cannot be measured in physiological low intracellular [Cl^−^], as they overlap with spike afterhyperpolarizations (AHPs). However, autaptic transmission was shown to modulate AHP duration (Pawelzik et al., 2003) and inter-spike intervals (Bacci and Huguenard, 2006) of cortical fast-spiking interneurons.

In control conditions (with intracellular 1 mM EGTA), PV cells showed a broad range of AHP durations (Fig. 6A,B), consistent with varying strengths of autaptic transmission across different PV cells (Pawelzik et al., 2003). When autaptic neurotransmission was blocked by intracellular perfusion of BAPTA (Fig. 4) (Bacci et al., 2003;Manseau et al., 2010), AHP duration was significantly smaller (17.96 ± 1.02 vs. 5.66 ± 0.42 ms; EGTA vs. BAPTA, respectively; p = 7.75E-16; independent *t-*test; Fig. 6A,B) and with a much reduced dispersion between cells (coefficient of dispersion: 2.6 vs. 0.9, EGTA vs. BAPTA, respectively, p = 4.06E-8; two-sample test for variance; Fig. 6B). This BAPTA-induced shortening of AHP likely resulted from the combined effect of this fast Ca^2+^ chelator on both autaptic transmission and Ca^2+^-activated K^+^ conductances, that are known to shape the AHP waveform (Sah and Faber, 2002).

**Figure 6.**
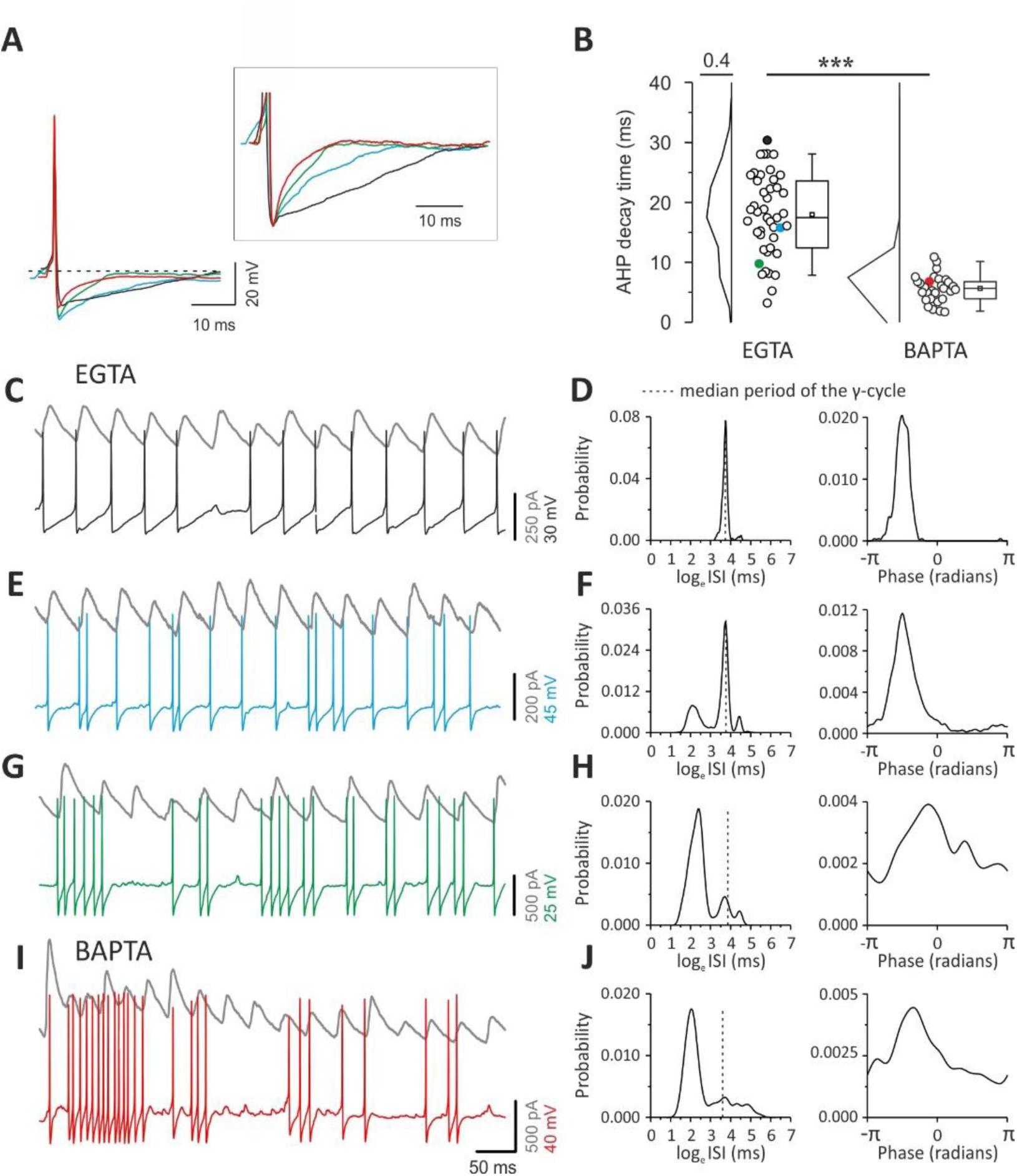
Different AHP durations and firing patterns of PV cells during γ-oscillations. ***A***, Representative overlapped action potentials (aligned to their peaks) recorded from different PV cells showing different AHP waveform in control (EGTA: black, blue and green traces) and in the presence of intracellular BAPTA (red trace). Inset: same traces at a larger voltage and time scale, normalized to the negative peak of the AHP. ***B***, Plots of AHP duration from PV cells recorded with control intracellular solution (EGTA, left) and BAPTA (right). Colors indicate the cells illustrated in A. ***C***, Representative traces of oscillating IPSCs recorded from a Layer II/III ChR2-negative PN (gray) and a PV cell with slow AHP (same of A-B, black). Note that spikes occur regularly at precise times, relative to the oscillating IPSCs. ***D***, Distributions of inter-spike intervals (ISIs, left) and phases (right) of the PV cell illustrated in C with a black trace. The dotted line indicates the interval corresponding to the median oscillation period. Note the sharp ISI distribution peaking at the oscillation period, and the sharp phase distribution. ***E-F***, Same as C-D, but for the PV cell represented with the blue trace in A-B. Note the appearance of spike doublets (E), yielding multimodal ISI distribution (F, left) and broader phase histogram (right). ***G-H***, Same as C-F, but for the cell represented with green trace in A-B. Note the appearance of high frequency bursts (G) yielding a large peak in the ISI distribution at faster intervals than the oscillation period. Further, note that the phase histogram yielded an even broader profile. ***I-J***, Same as in C-H but for the PV cell intracellularly perfused with BAPTA, illustrated with a red trace in A-B. Note the similar firing behavior of the EGTA cell characterized by the fast AHP and burst firing (green traces in A, B, G, H).

Under control (EGTA) conditions, the specific duration of the AHP determined the coupling of PV-cell spikes with γ-oscillations. Indeed, PV cells with slow AHPs, produced spike trains, which were regular and strongly coupled to γ-oscillations, as the large majority of action potentials occurred almost invariably at a precise time during the γ-cycle (Fig. 6C). This strong coupling of PV-cell spiking activity with γ-oscillations determined a very sharp, unimodal distribution of inter-spike intervals (ISIs) peaking at the γ-oscillation period, as well as a sharp phase coupling histogram (Fig. 6D). PV cells with faster AHPs exhibited high-frequency doublets (Fig. 6E,F), and/or bursts of spikes (Fig. 6G,H). In these cases, ISI distributions were multimodal, exhibiting peaks at shorter intervals than the oscillation period. Moreover, the spike coupling to γ-phase was increasingly less sharply distributed (Fig. 6F,H).

Multi-modality of ISI distributions and broad spike-phase coupling resulted from an increasing number of spikes with very fast intervening ISIs, not effectively matching the period of ongoing γ-rhythm in Layer II/III (Fig. 6F,H). Control PV cells characterized by the shortest AHPs and consequent weak coupling with γ-oscillations (Fig. 6G,H) exhibited firing patterns that were similar to PV cells, in which autapses were blocked by intracellular perfusion of BAPTA, and characterized by fast AHPs.

These PV cells intracellularly perfused with BAPTA consistently produced high frequency bursts of spikes (Fig. 6I), yielding ISI distributions with the largest peak at a faster interval than the oscillation period and broad spike-phase coupling distributions (Fig. 6J).

On average, the distribution of ISIs in control (EGTA) cells peaked close to the γ-cycle. Conversely, BAPTA-filled PV interneurons discharged with ISIs not matching the γ-period (mean log ratio: −0.147 ± 0.077 and −1.078 ± 0.155 in EGTA and BAPTA, respectively, p= 6.45E-7, Wilcoxon rank-sum; Fig. 7A). Further, distributions of ISIs were significantly less dispersed in control (EGTA) PV cells as compared to PV cells filled with BAPTA as measured by the log ISI entropy (mean: 6.51±0.08 and 6.87±0.11 bits in EGTA and BAPTA, respectively, p= 0.0063, Mann-Whitney; Fig. 7A). For each PV cell, the slower the AHP, the closer to the γ-period was its ISI, whereas PV cells exhibiting fast AHP (such as those whose autapses were blocked by intracellular BAPTA) fired with ISIs that were faster than the γ-period (Spearman R = 0.553; p = 4.02E-7; Fig. 7B).

**Figure 7.**
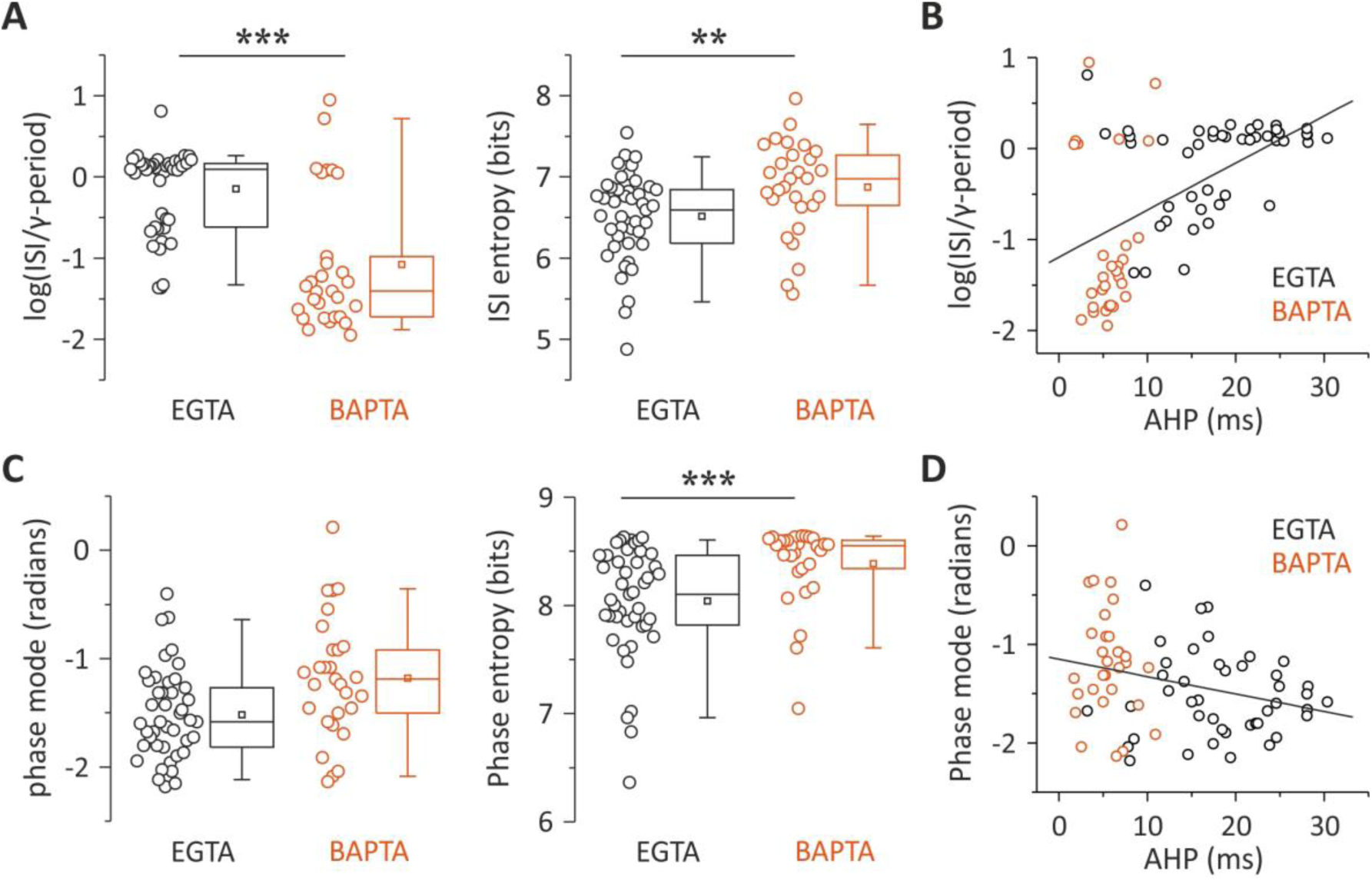
Autaptic neurotransmission lock PV-cell firing to γ-oscillations. ***A***, Population data and distributions of ISI values relative to γ-period (left) and ISI entropy (right) of PV cells recorded with intracellular EGTA (black) and BAPTA (red). The zero value in the Y-axis of the left plot corresponds to the period of the γ-cycles. ***B***, AHP durations plotted as a function of ISI values relative to γ-period for PV cells recorded in EGTA (black) and BAPTA (red). Black line: linear regression. ***C-D***, Same as in A-B but for the peak (mode) of phase distributions as illustrated in Fig. 6. ** p<0.01; *** p<0.001.

The peaks of phase distributions were not significantly different in EGTA and BAPTA (−1.524 ± 0.0613 and −1.189 ± 0.098 radians respectively, p=0.1, k-test for circular distribution Fig. 7C). However the dispersion of their distributions were different in control (EGTA) vs. BAPTA-filled PV cells, indicating a lesser extent of phase lock induced by intracellular perfusion of BAPTA as measured by the entropy of the phase distribution (mean: 8.041±0.078 and 8.387±0.0689 bits in EGTA and BAPTA, respectively, p= 4.6E-4,; Fig. 7C). Also in the case of phase, a significant correlation was found between AHP duration and phase (Spearman R = −0.285; p = 0.0145; Fig. 7D).

Altogether, these results suggest that the strength of autaptic self-inhibition determines the coupling of PV-cell spiking to γ-oscillations, by modulating the duration of their own AHPs.

## Discussion

Here we found that autaptic transmission is overall the most powerful output from PV cells in neocortical Layer V. Autaptic transmission is ~3-fold stronger than synaptic inhibition onto PNs, and ~2-fold larger than PV-PV connections. Moreover, we found that PV cells with strong autaptic transmission produce a weaker synaptic output onto other PV cells and vice versa, thus defining a novel architecture of relative connectivity strength. Despite the existence of a minority of PV cells with stronger PV-PV synaptic than autaptic transmission, self-inhibitory autapses, originating from a single axon, contribute up to ~40% of the entire perisomatic inhibition onto PV cells. Strong, reliable and fast autaptic self-inhibition of PV cells contributes to duration of the AHP and therefore the ISIs of these interneurons, affecting their degree of synchronization with γ-oscillations.

The observation of larger autaptic currents than inhibitory synaptic responses elicited by the same PV cells onto PNs was not due to differences in the number of release sites, but to a larger autaptic quantal size. A larger quantal size can be ascribed to several causes, including, for example, different subunit composition of GABA_A_Rs, their expression level at postsynaptic sites, their phosphorylation state, and the specific interactions with distinct scaffolding, anchoring and trans-synaptic proteins (Fritschy et al., 2012).

Another reason for a smaller quantal size in PNs could be a more distal location of PV-PN synapses as opposed to PV cell autapses. This could result in more low-pass filtering of synaptic responses with a consequent reduction in their size. Although we cannot exclude that this is the case, we argue against this possibility, since PV cells are known to be perisomatic targeting (Freund and Katona, 2007). Indeed, the cell body of large Layer V PNs is almost completely innervated by PV-positive inhibitory terminals (Bodor et al., 2005). This is consistent with the very fast rise-time of PV-PN unitary synaptic responses (<1 ms, data not shown). Finally, although a different quantal size between two synaptic connections is traditionally ascribed to postsynaptic factors, we cannot exclude that the difference in q could be due to a different amount of neurotransmitter released by each vesicle at each individual synapse. Future studies will be necessary to pinpoint the molecular mechanism underlying the difference in quantal size between autapses onto PV cells and synapses onto PNs.

Curiously, connections between PV cells showed a connectivity logic dictated by their actual autaptic strength. Although self-contacts were generally stronger than heterosynaptic connections with other PV cells, autapses were weaker in a minority of cases (~38%). In both ‘introverted’ and ‘extroverted’ PV cells, the difference between autaptic and synaptic strength was due to a higher or lower number of release sites, and thus it was due to anatomical specificities. Similar quantal size at autaptic and synaptic connections between PV cells indicates that postsynaptic sensitivity to released GABA at autaptic contacts is equivalent to that of synaptic connections. This could be due to expression of molecularly similar postsynaptic receptor clusters, and similar degree of autaptic and synaptic filtering.

The existence of ‘extroverted’ and ‘introverted’ PV cells prompts the question of whether they belong to different cell types. Whereas we detected no changes of passive and firing properties of ‘introverted’ and ‘extroverted’ (data not shown), we cannot exclude differential morphology and/or connectivity patterns. Alternatively, the differential strength of self-vs. heterosynaptic inhibitory contacts could be due to activity-dependent plasticity of GABAergic connections from PV cells. Indeed, it has been shown that postsynaptic activity could modulate the strength of GABAergic synapses from PV cells in the visual (Xue et al., 2014) and somatosensory cortex (Lourenco et al., 2014). Future studies will be necessary to reveal the mechanisms underlying the differential autaptic and synaptic strength onto PV cells.

Functional autaptic neurotransmission represents a powerful form of fast disinhibition of PV cells. Accumulating evidence indicates that disinhibitory circuits play crucial roles for several cognitive functions (Kepecs and Fishell, 2014;Letzkus et al., 2015;Pi et al., 2013;Tremblay et al., 2016). In particular, disinhibition operated by VIP cells (Gulyas et al., 1996;Pfeffer et al., 2013) may be crucial for auditory discrimination (Pi et al., 2013), memory retention in prefrontal cortex (Kamigaki and Dan, 2017) and other forms of associative learning and memory (Letzkus et al., 2015). Importantly, however, VIP cell-mediated disinhibition requires multi-synaptic circuits, and, because it occurs over relatively long (100s of ms) time windows, it might be important for modulating the information carried by a whole spike train according to a traditional rate-coding scheme. By contrast, autaptic self-inhibition of PV cells accounts for ~40% of the total perisomatic inhibition they received. It is fast (occurring at a millisecond timescale) and activated by single spikes. Autaptic disinhibition of PV cells should therefore be crucial for encoding information carried by the precise timing of individual spikes within a high-frequency train. Indeed, we show that fast GABAergic self-inhibition of PV cells modulates the locking of their spike timing to network oscillations in the β-γ-frequency range.

Autaptic transmission occurs immediately after single action potentials, thus modulating the duration of the AHPs of PV cells during trains of spikes (Bacci and Huguenard, 2006;Pawelzik et al., 2003). In control (EGTA) conditions, we found a broad range of AHP durations. This is consistent with heterogeneous autaptic strengths among several PV cells, and lack of functional self-innervation in some cases (~30%). Accordingly, autaptic blockade by intracellular BAPTA invariably produced fast AHPs and high frequency firing. Relatively long-lasting AHPs correlated with a strong synchronization of PV-cells output with γ-oscillations. Importantly, the tight locking of PV-cell spikes to γ-activity shown here was similar to that recorded from PV cells in the visual cortex *in vivo* in the absence and presence of sensory stimuli (Perrenoud et al., 2016).

Interestingly, faster AHPs were responsible for the generation of high frequency doublets and/or bursts of spikes. This activity could be detected in virtually all cells intracellularly perfused with BAPTA. The sharpening of PV-cell AHPs induced by intracellular BAPTA was likely due to the blockade of autaptic transmission combined to the impairment of Ca^2+^-activated K^+^ channels, both contributing to AHP peak and duration (Sah and Faber, 2002). Interestingly, a minority of cells recorded with intracellular control (EGTA) conditions exhibited sharp AHPs similar to those recorded with intracellular BAPTA. This is consistent with the fraction of PV interneurons lacking functional autaptic transmission, but with intact Ca^2+^-activated K^+^ channels. The heterogeneity of AHP durations and firing behaviors during γ-activity in control PV cells suggests that the instantaneous spike frequency is highly controlled by autaptic strength. A strong GABAergic conductance, reliably activated with a high release probability immediately after each spike, shapes the window of opportunity to fire a subsequent spike. Therefore, rhythmic glutamatergic activation of PV cells by Layer II/III PNs and the strong, fast and reliable autaptic self-inhibition work in synergy to lock PV-cell firing to the oscillation period. Given the crucial role of mutual inhibition between PV cells during synchronous network activity (Cardin et al., 2009;Sohal et al., 2009), spike timing regulation through autaptic self-inhibition will thus strongly influence the output spike timing of several PV cells in a millisecond timescale effectively synchronizing networks of PV cells during the emergence of fast oscillations.

Overall, our results indicate that self-inhibition of PV cells via autaptic neurotransmission is among the most powerful connections from this cell type within the layer 5 cortical microcircuit promoting their spiking synchronization during γ-oscillations. GABAergic autaptic self-inhibition of PV cells is therefore an important mechanism underlying the key role of these cells during cognitive-relevant network oscillations, with possible crucial consequences in both physiological and pathological cortical operations.

## Methods

### Animals

Experimental procedures followed national and European (2010/63/EU) guidelines and were approved by the authors’ institutional review boards and national authorities. All efforts were made to minimize suffering and reduce the number of animals. Experiments were performed on C57BL/6J mice obtained by breeding PV-cre mice to a reporter line harboring a loxP-flanked STOP cassette associated to the red fluorescent protein variant tdTomato (line Ai14 jax line 007914). This mouse line allowed recognition of PV interneurons in live acute slices. Indeed, tdTomato-expressing cells had typical multipolar morphology, aspiny dendrites and fast-spiking behavior (not shown).

### In utero electroporation

Timed-pregnant PV-cre female mice bred with tdTomato males (15.5 days postcoitum) were anaesthetized with 1-2% isoflurane. The abdomen was cleaned with 70% ethanol and swabbed with betadine. Buprenorphine (0.05 mg/kg) was administered subcutaneously for preoperative analgesia and local anesthetic bupivacaine (2.5mg/kg) was injected between the skin and the abdomen 5 min before incision. A midline ventral laparotomy was performed and the uterus gently exposed and moistened with PBS pre-warmed at 37 °C. pCAG-mRFP (0.8 μg/μl) (Addgene #28311) (Manent et al., 2009) (plasmid DNA was mixed with pCAG-ChR2-Venus (0.8 μg/μl) (Addgene #15753) (Petreanu et al., 2007) and Fast Green (0.025%; Sigma) in saline solution (PBS).

Each embryo was injected with the mix DNA solution through the uterine wall into the lateral ventricle using pressure-controlled bevelled glass capillaries (WPI micropipette beveler). After each injection, tweezers disk electrodes (platinium 5mm round, Sonidel) were positioned at 0° angle with respect to the rostral-caudal axis of the head of each embryos and voltage pulses (5 pulses, 40 V; 50 ms; 5Hz) were applied to electroporate the DNA (square wave electroporator, Nepa Gene). The uterine horn containing the embryos was then placed back into the peritoneal cavity and moistened with PBS. The abdomen and skin were sutured and the latter was cleaned with betadine. The procedure typically lasted maximum 40 min starting from anesthesia induction. Pups were born by natural birth and screened for location and strength of transfection by trans-cranial epifluorescence under a fluorescence stereoscope.

### In vitro slice preparation

Naïve or *in utero* electroporated mice were deeply anesthetized with isoflurane, decapitated and the brains quickly removed. Coronal slices were prepared from somatosensory cortex of mice aged (P15-P25) using a vibratome (Leica VT1200 S) in a free or reduced sodium cutting solution (4°C). Slices were initially stored at 34°C for 30 min in standard or reduced sodium solution (ASCF) then at room temperature for at least 1h before being transferred to a submerged recording chamber maintained at ~30°C.

For unitary autaptic and synaptic IPSCs experiments, coronal slices (350 μm thick) were obtained from somatosensory cortex using a free sodium cutting solution containing (in mM): choline 118, glucose 16, NaHCO_3_, 26, KCl 2.5, NaH_2_PO_4_ 1.25, MgSO_4_ 7, CaCl_2_ 0.5, pyruvic acid 3, myo-inositol 3, ascorbic acid 0.4 gassed with 95% O_2_ and 5% CO_2_. Then, slices were stored in oxygenated standard ASCF (in mM): NaCl 126, KCl 2.5, CaCl_2_ 2, MgSO_4_ 1, NaH_2_PO_4_ 1.25, NaHCO_3_ 26, glucose 20; pH 7.4. For photo-induced gamma oscillations experiments, 400 μm-thick coronal somatosensory cortical slices were prepared from the transfected hemisphere. Slices were cut and stored in oxygenated reduced sodium ACSF containing (in mM): NaCl 83, sucrose 72, glucose 22, NaHCO_3_ 26, KCl 2.5, NaH_2_PO_4_ 1, MgSO_4_ 3.3, CaCl_2_ 0.5, pyruvic acid 3, myo-inositol 3, ascorbic acid 0.4;pH 7.4.

### Electrophysiology

#### Unitary autaptic and synaptic IPSCs

Recordings were obtained in standard ACSF at 30°C from PV-PV cells pairs and PV-PN pairs of Layer V primary barrel somatosensory cortex. Neuron types were visually determined using infrared video microscopy. PV interneurons were visible as tdTomato positive fluorescent cells whereas PNs were identified by their large soma and emerging apical dendrite together with firing behavior. Whole-cell voltage-clamp recordings were obtained with patch pipettes (2-4 MΩ) filled with a high [Cl-] intracellular solution containing (in mM): K-gluconate 70, KCl 70, Hepes 10, EGTA 1, MgCl_2_ 2, MgATP 4, NaGTP 0.3 or K-gluconate 35, KCl 70, Hepes 10, 4K-BAPTA 20, CaCl_2_ 2, MgATP 4, NaGTP 0.3; pH 7.2 adjusted with KOH; 290 mOsm; for EGTA and BAPTA experiments, respectively. GABA_A_ receptor-mediated IPSCs were isolated by adding 6,7-dinitroquinoxaline-2,3,dione (DNQX, 10 μM) in the bath perfusion and recorded at a holding potential of –80 mV or −70 mV. For miniature IPSCs (mIPSCs) recordings, DNQX and tetrodotoxin (TTX, 1 μM) were added to the bath perfusion. When indicated, SR95531 [6-imino-3-(4-methoxyphenyl)-1(6H)-pyridazine-butanoic acid hydribromide] (gabazine, 10 μM) was also applied by bath perfusion to block GABAA receptors. All drugs were obtained from Tocris Cookson (Bristol, UK).

#### Photo-induced γ-oscillations

Once being transferred to the submerged recording chamber, slices were superfused with modified ACSF containing (in mM): NaCl 119, KCl 2.5, CaCl_2_ 2,5, MgSO_4_ 1,3, NaH_2_PO_4_ 1.3, NaHCO_3_ 26, glucose 20 (pH 7.4) maintained at 30°C. Before starting recordings, slices were carefully examined to check mRFP expression in Layer II/III of the somatosensory cortex. Whole-cell, voltage-clamp recordings of photo-induced γ-oscillations were obtained from ChR2-negative PNs identified by the pyramidal shape of their soma, the emerging apical dendrite and the absence of mRFP- and tdTomato fluorescence. Patch pipettes were filled with a cesium-based low [Cl-] intracellular solution containing (in mM): CsMeSO4^−^ 125, CsCl 3, Hepes 10, EGTA 5, MgCl_2_ 2; MgATP 4, NaGTP 0.3, QX314-Cl 5; pH 7.2 corrected with CsOH; 290 mOsm. Inhibitory (IPSCs) and/or excitatory (EPSCs) postsynaptic currents were recorded at GluR and GABA_A_R reversal potentials, respectively. Simultaneous current-clamp recording were obtained from layer 5 tdTomato-positive fluorescent PV cells located within the same cortical column of the layer 2/3 PN, using a Kgluconate-based low [Cl-] intracellular solution containing (in mM): K-gluconate 120, KCl 13, Hepes 10, EGTA 1, MgCl_2_ 2, MgATP 4, NaGTP 0.3 or K-gluconate 103, KCl 13, Hepes 10, 4K-BAPTA 20, CaCl_2_ 2, MgATP 4, NaGTP 0.3; pH 7.2 adjusted with KOH; 290 mOsm; for EGTA and BAPTA conditions, respectively.

### Photo-stimulation

Photo-stimulation was induced using a blue LED (λ = 470 nm, OptoLED Lite, Cairn research, UK) collimated and coupled to the epifluorescence path of the microscope (BX51WI; Olympus). Light was delivered through a 60X (1.0 NA) water immersion lens, centered on Layer II/III. The light intensity and waveform was controlled by the analog output of a digitizer (Digidata 1440A, Molecular Devices). Light ramps had a duration of 1-3 s, a slope of 0.1-0.8 mW s^-1^, started at zero intensity and reached a final intensity of 0.3-1.6 mW s^-1^. The slope was adjusted in each slice to obtain a robust rhythmic activity in the gamma frequency range with a stable power for the entire duration of the stimulus. Light ramps were repeated with a 60 s interval.

### Data acquisition and analysis

Signals were amplified using a Multiclamp 700B patch-clamp amplifier (Molecular Devices, USA), sampled at 20 KHz and filtered at 2 KHz or 10 KHz in voltage-clamp and current-clamp mode, respectively. Voltage measurements were not corrected for liquid junction potential. Access resistance was <20 MΩ and monitored throughout the experiment. Recordings were discarded from analysis if the resistance changed by >20% over the course of the experiment. Data were analyzed using pClamp (Molecular devices, USA), Origin (Microcal Inc., USA), MATLAB (MathWorks, USA) and custom written scripts and software.

#### Unitary autaptic and synaptic IPSCs

A brief (0.2-0.6 ms) depolarizing current step was injected in the presynaptic PV interneuron from PV-PV or PV-PN pairs. Voltage jumps were calibrated for each stimulated PV cell to a value ranging between −20 and 0 mV from the holding potential, to reduce the contribution of K^+^-mediated conductance of the action current, contaminating the autaptic response. GABAergic autaptic and synaptic responses were recorded in the stimulated PV cell itself and in the paired cell (PV or PN), respectively, in whole-cell voltage-clamp mode. For quantal parameters analysis, responses were recorded at two extracellular Ca^2+^ concentrations (1.5 and 2.0 mM) to induce low and high release probabilities, respectively. Gabazine was applied at the end of the recordings to subtract the stimulus waveform and the isolated action current to autaptic responses (Fig 1A,C). The IPSCs amplitude was estimated as the current from the baseline before the onset of the stimulus to the peak on control or subtracted trace when gabazine was applied. Data were analyzed using pClamp (Molecular devices, USA), Origin (Microcal Inc., USA) and MATLAB (Mathworks, USA) software.

#### Bayesian quantal analysis

Quantal parameters were estimated using an improved implementation of Bayesian Quantal Analysis (Bhumbra and Beato, 2013) as described previously (Bhumbra et al., 2014;Moore et al., 2015). Briefly, synaptic or autaptic currents were measured in the presence of two different Ca^2+^ concentrations (1.5 and 2 mM) corresponding to intermediate and high release probabilities. BQA was performed only for experiments in which at least 50 stable responses per condition could be recorded, followed (in the case of autaptic connections) by application of gabazine in the presence of both Ca^2+^ concentrations, in order to subtract the profile of the action currents, that is subject to changes following reduction of Ca^2+^.

In contrast to multiple probability fluctuation analysis (MPFA) (Silver, 2003) that relies on parabolic fits to the variance-mean relationship of synaptic currents, BQA models the distribution of all amplitudes observed at different release probabilities. The advantage this confers on BQA is that quantal parameters can be reliably estimated from few response measurements obtained from only two different release probabilities (Bhumbra and Beato, 2013). Quantal parameters were estimated as the median value of the posterior distributions. The BQA implementation was modified in the selection of the marginal priors for the number of release sites. Contrary to our previous implementation (Bhumbra and Beato 2013), in which the marginal priors for the probability of release and the uniquantal coefficient of variation were assigned according to Jeffrey’s rule, while the number of release sites had an uniform prior, here we applied Jeffrey’s rule to the number of release sites as well, resulting in a reciprocal, rather than uniform, prior distribution.

#### Miniature inhibitory synaptic events

ACSF containing a high concentration of K^+^ (~20 mM) was applied using a pressure system (puff), through a glass pipette located near the cell body of the recorded PV interneuron to depolarize axon terminals impinging the recorded neuron. High-K^+^ puffs induced global asynchronous release of GABA that could be detected as a substantial increase in the frequency of miniature inhibitory postsynaptic currents (mIPSCs). mIPSCs were recorded during successive sequences of baseline activity and throughout puff application (3-6 s, 1 puff / minute), for at least 20 min. Miniature GABAergic events were detected using a custom written software (Detector, courtesy J.R. Huguenard, Stanford University; Supplemental figure 1). Briefly, individual events were detected with a threshold-triggered process from a differentiated copy of the raw current trace. Detection frames were inspected visually to ensure that the detector was working properly. mIPSC frequency was calculated for successive 1 s time windows. For each puff application, the relative mIPSC frequency was estimated as the ratio between the maximum mean frequency (1 s bin) during puff application and the average of the mean frequencies for the entire baseline duration. To evaluate the percentage of mIPSC frequency decrease, we compared the relative frequency (average of 3 successive values) at the beginning of the recording (5-10 min after whole-cell configuration establishment) to the relative frequency after the block of autaptic currents by BAPTA (~20 min after whole-cell configuration establishment).

#### Firing properties of layer 5 PV cells during photo-induced γ-activity

Bursts of γ activity were evoked by light stimulation of ChR2-positive PN cell bodies in Layer II/III. While recording from a layer 5 PV interneuron, a simultaneous recording of a Layer II/III ChR2-negative PN was used to determine the period of the γ activity. Rhythmic synaptic events evoked by light stimulation were detected using *Detector (*courtesy of J.R. Huguenard) as described above for mIPSCs. Spikes of Layer V PV cells were extracted using a threshold of −10 mV on the membrane potential trace. PSC cycles were measured and the timing of each spike in the PV neuron was expressed as a phase relative to the peak of each γ oscillation. Inter-spike intervals (ISI) and phase distributions were computed for each cell using custom written software (MATLAB). Variability of firing was evaluated using the entropy of the log interval distribution (Bhumbra and Dyball, 2004, 2010), whereas the dispersion of peri-cycle spike times was quantified using the entropy of the corresponding phase distribution (Bhumbra and Dyball, 2010).

The Circular statistic toolbox of MATLAB was used to compute parameters of phase distributions and their associated statistical tests, as indicated in the text.

*Afterhyperpolarization (AHP) duration of single action potentials* was measured as the 10-90% decay time setting the baseline right before the spike (5 ms window). Hence, isolated spikes were selected for this analysis since the decay time of action potential being part of doublets or burst could be contaminated by the generation of the following one. The average value of the AHP decay time of 10-20 spikes was considered for each PV cell.

#### Statistics

Since most data distributions were not normal, unless indicated in the text, we used non-parametric significance test, Wilcoxon’s signed-rank test and Wilcoxon’s rank-sum test for paired and unpaired data respectively.

## Author Contributions

CD and AB conceived the project; CD performed all the recordings and analyzed the data; GSB and MB designed and performed the quantal analysis; GSB, MB, AP designed and performed the analysis on γ-oscillations; CD, CM and AA performed *in utero* electroporations; CD, GSB, MB and AB wrote the paper; MB and AB supervised the project.

## Acknowledgments

We thank Frédéric Manseau, Pasqualina Farisello and Tommaso Fellin for their initial involvement in this project and Geeske M. van Woerden for help with *in utero* electroporation. We are grateful to Joana Lourenço, Javier Zorrilla de San Martin, Nelson Rebola and David DiGregorio for critically reading this manuscript. This work was supported by European Research Council (ERC) under the European Community’s 7th Framework Programmme (FP7/2007-2013)/ERC grant agreement No 200808); “Investissements d’avenir” ANR-10-IAIHU-06; Agence Nationale de la Recherche (ANR-13-BSV4-0015-01, ANR-FRONTELS and ANR-NanoSynDiv), Fondation Recherche Médicale (Equipe FRM DEQ20150331684), NARSAD independent investigator grant, and a grant from the Institut du Cerveau et de la Moelle épinière (Paris) (A.B.) and by a Leverhulme Trust Research grant (RPG-2013-176) and a Biotechnology and Biological Sciences Research Council Grant (BB/L001454) to M.B.

## Supplemental Information

**Supplemental Figure 1: Detection of global inhibition onto PV cells induced by ambient depolarization by high extracellular K**^**+**^.

Global inhibition onto single PV cells was estimated as the increase of mIPSC frequency evoked by a local puff of 20 mM KCl, triggering massive Ca^2+^-dependent release of GABA onto the recorded neuron (Fig. 4). Shown is a snapshot of the mIPSC detection software before (left) and after (right) the high KCl puff, illustrating the ability of detecting high-frequency synaptic events in response to ambient depolarization. Events were detected based on a threshold-crossing algorithm on the derivative (bottom) of the current traces (top). Vertical lines indicate detected synaptic events.

## Supplemental Information

**Supplemental Figure 1:**
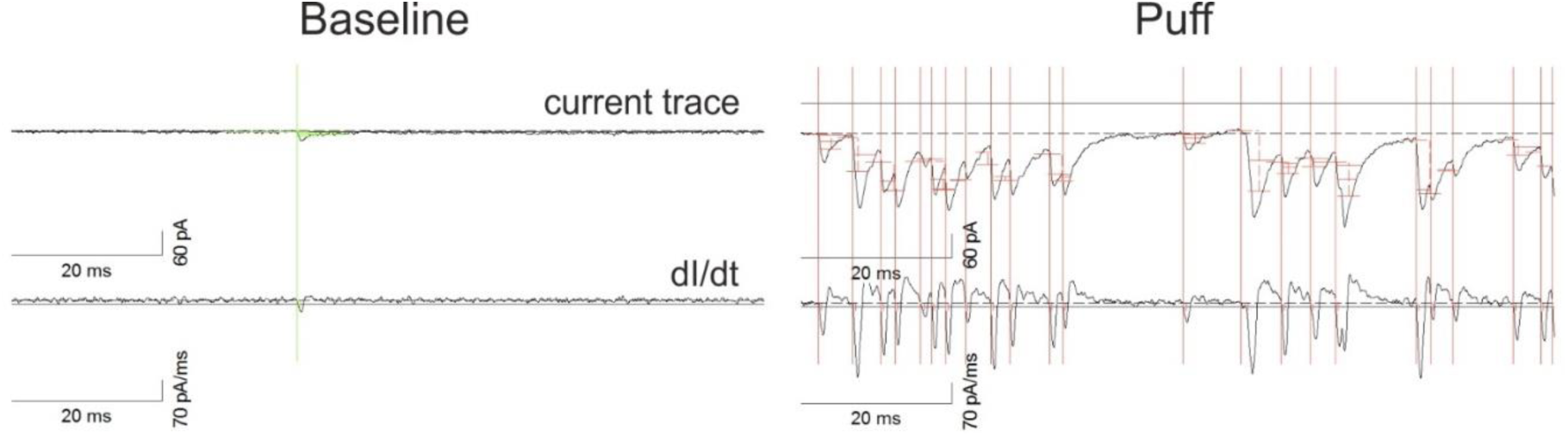
Detection of global inhibition onto PV cells induced by ambient depolarization by high extracellular K^**+**^. Global inhibition onto single PV cells was estimated as the increase of mIPSC frequency evoked by a local puff of 20 mM KCl, triggering massive Ca^2+^-dependent release of GABA onto the recorded neuron (Fig. 4). Shown is a snapshot of the mIPSC detection software before (left) and after (right) the high KCl puff, illustrating the ability of detecting high-frequency synaptic events in response to ambient depolarization. Events were detected based on a threshold-crossing algorithm on the derivative (bottom) of the current traces (top). Vertical lines indicate detected synaptic events.

